# Enrichment of rare protein truncating variants in amyotrophic lateral sclerosis patients

**DOI:** 10.1101/307835

**Authors:** Sali M.K. Farhan, Daniel P. Howrigan, Liam E. Abbott, Andrea E. Byrnes, Claire Churchhouse, Hemali Phatnani, Bradley N. Smith, Simon D. Topp, Evadnie Rampersaud, Gang Wu, Joanne Wuu, Amelie Gubitz, Joseph R. Klim, Daniel A. Mordes, Sulagna Ghosh, CReATe Consortium, FALS Consortium, ALSGENS Consortium, Kevin Eggan, Rosa Rademakers, Jacob L. McCauley, Rebecca Schüle, Stephan Züchner, Michael Benatar, J. Paul Taylor, Mike A. Nalls, Bryan Traynor, Christopher E. Shaw, David B. Goldstein, Matthew B. Harms, Mark J. Daly, Benjamin M. Neale

**Affiliations:** Analytic and Translational Genetics Unit, Department of Medicine, Massachusetts General Hospital and Harvard Medical School, Boston, MA 02114, USA.; Program in Medical and Population Genetics, Broad Institute of Harvard and MIT, 7 Cambridge Center, Cambridge, MA 02142, USA.; Stanley Center for Psychiatric Research, Broad Institute of Harvard and MIT, Cambridge, MA 02142, USA.; Center for Genomics of Neurodegenerative Disease, New York Genome Center, New York, NY 10013, USA.; United Kingdom Dementia Research Institute Centre, Maurice Wohl Clinical Neuroscience Institute, Institute of Psychiatry, Psychology and Neuroscience, King’s College London, London, SE5 9NU, U.K.; Department of Computational Biology, St. Jude Children’s Research Hospital, Memphis, TN 38105, USA.; Department of Neurology, University of Miami, Miami, FL 33136, USA.; National Institute of Neurological Disorders and Stroke, National Institutes of Health, Bethesda, MD 20824, USA.; Department of Stem Cell and Regenerative Biology, Harvard Stem Cell Institute, Harvard University, Cambridge, MA 02138, USA.; Department of Neuroscience, Mayo Clinic, Jacksonville, FL 32224, USA.; John P. Hussman Institute for Human Genomics, University of Miami, Miller School of Medicine, Miami, FL 33136, USA.; The Dr. John T. Macdonald Foundation Department of Human Genetics, University of Miami, Miller School of Medicine, Miami, FL 33136, USA.; Center for Neurology and Hertie Institute für Clinical Brain Research, University of Tübingen, Tübingen 72076, Germany. German Center for Neurodegenerative Diseases (DZNE), Tübingen 72076, Germany.; Howard Hughes Medical Institute, Chevy Chase, MD 20815, USA.; Department of Cell and Molecular Biology, St. Jude Children’s Research Hospital, Memphis, TN 38105, USA.; Molecular Genetics Section, Laboratory of Neurogenetics, National Institute on Aging, Bethesda, MD 20892, USA.; Data Tecnica International, Glen Echo, MD 20812, USA.; Neuromuscular Diseases Research Section, Laboratory of Neurogenetics, National Institute on Aging, NIH, Porter Neuroscience Research Center, Bethesda, MD 20892, USA.; Department of Neurology, Johns Hopkins University, Baltimore, MD 21287, USA.; Centre for Brain Research, University of Auckland, Auckland, New Zealand.; Institute for Genomic Medicine, Columbia University, New York, NY 10032, USA.; Department of Neurology, Columbia University, New York, NY 10032, USA.

**Keywords:** Amyotrophic lateral sclerosis, protein truncating variants, neurodegeneration, rare variant associations, DNAJC7

## Abstract

To discover novel genetic risk factors underlying amyotrophic lateral sclerosis (ALS), we aggregated exomes from 3,864 cases and 7,839 ancestry matched controls. We observed a significant excess of ultra-rare and rare protein-truncating variants (PTV) among ALS cases, which was primarily concentrated in constrained genes; however, a significant enrichment in PTVs does persist in the remaining exome. Through gene level analyses, known ALS genes, *SOD1, NEK1*, and *FUS*, were the most strongly associated with disease status. We also observed suggestive statistical evidence for multiple novel genes including *DNAJC7*, which is a highly constrained gene and a member of the heat shock protein family (HSP40). HSP40 proteins, along with HSP70 proteins, facilitate protein homeostasis, such as folding of newly synthesized polypeptides, and clearance of degraded proteins. When these processes are not regulated, misfolding and accumulation of degraded proteins can occur leading to aberrant protein aggregation, one of the pathological hallmarks of neurodegeneration.

## INTRODUCTION

Amyotrophic lateral sclerosis (ALS) is a late-onset neurodegenerative disease characterized primarily by degeneration of motor neurons leading to progressive weakness of limb, bulbar, and respiratory muscles (Al-Chalabi et al., 2017; Strong et al., 2017). Genetic variation is an important risk factor for ALS. Given that 5-10% of patients report a positive family history (Al-Chalabi et al., 2017) and ~10% of sporadic patients carry known familial ALS gene mutations, the distinction between familial and sporadic disease is increasingly blurred (Al-Chalabi, 2017). Until recently, ALS gene discoveries were made through large multigenerational pedigrees in which the gene and the causal variant segregated in an autosomal dominant inheritance pattern with very few cases of autosomal recessive inheritance reported. Collecting sporadic case samples has been valuable for gene discovery in more common disorders such as schizophrenia (Singh et al., 2016), inflammatory bowel disease (Mohanan et al., 2018), and type 2 diabetes (Manning et al., 2017), and can have profound effects on the success of targeted therapeutic approaches (Al-Chalabi et al., 2017; Hamburg and Collins, 2010; Nelson et al., 2015). The most recent ALS genetic discoveries using large massively parallel sequencing data yielded several gene discoveries, *TBK1, TUBA4A, ANXA11* and *NEK1* and *KIF5A* (Cirulli et al., 2015; Kenna et al., 2016; Nicolas et al., 2018; Smith et al., 2014; Smith et al., 2017); in addition to other risk loci in *C21orf2, MOBP*, and *SCFD1* (van Rheenen et al., 2016).

Herein, we have assembled the largest ALS exome case-control study to date, consisting of 11,703 individuals (3,864 cases and 7,839 controls). We complemented our analysis by leveraging allele frequencies from large external exome sequencing databases in DiscovEHR (>50,000 samples; cite DiscovEHR) and a subset of ExAC (>45,000 samples) (Lek et al., 2016). In our analysis, we observed an excess of rare protein truncating variants (PTV) in ALS cases, which primarily resided in constrained genes. There was no enrichment of rare PTVs in genes known to confer risk to ALS; genes associated with clinically overlapping diseases; or genes in which their expression is specific to the brain.

Furthermore, through gene burden testing, we confirmed the known association of *SOD1, NEK1*, and *FUS*, while also observing PTVs in *DNAJC7* in 4/3,864 cases and 0/28,910 controls. *DNAJC7* is a highly constrained gene, and encodes a DNAJ molecular chaperone, which facilitates protein maintenance and quality control, such as folding of newly synthesized polypeptides, and clearance of degraded proteins (Lackie et al., 2017). Dysregulation of these processes can lead to aberrant protein aggregation, one of the pathological hallmarks of neurodegenerative diseases.

## RESULTS

### Patient demographics and dataset overview

We processed our initial dataset of 15,722 samples through a rigorous quality control pipeline using Hail, an open-source, scalable framework for exploring and analyzing genomic data https://hail.is/. We removed samples with poor sequencing quality, high levels of sequence contamination, closely related with one another, ambiguous sex status, or population outliers per PCA (Table S1 and Table S3). Our final data set consisted of 11,703 samples: 3,864 cases and 7,839 controls. Individuals were of European descent with 7,355 samples (62.8%) classified as males. Of 3,864 cases, 2,274 samples (58.9%) were classified as males; where 5,081 samples (64.8%) were classified as males in controls.

### Excess of exome-wide rare protein truncating variants

Using multiple firth based logistic regression analyses incorporating sample sex, PC1-PC10, and the total exome count (summation of synonymous variants, benign missense variants, damaging missense variants, and PTVs), we observed a significant enrichment of singleton PTVs in ALS cases relative to controls (OR: 1.07; 95% CI: 1.04-1.10; P-value: 5.00×10^-^7); ultra-rare singleton PTVs (OR: 1.08; 95% CI: 1.05-1.11; P-value: 1.97×10^-^6); and rare PTVs (OR: 1.04; 95% CI:1.02-1.06; P-value: 1.77×10^-^7) (Figure 1). These values all passed multiple test correction (P<0.0125). The number of doubletons (AC=2) was too low to be sufficiently powered to detect any significant enrichment.

**Figure 1.**
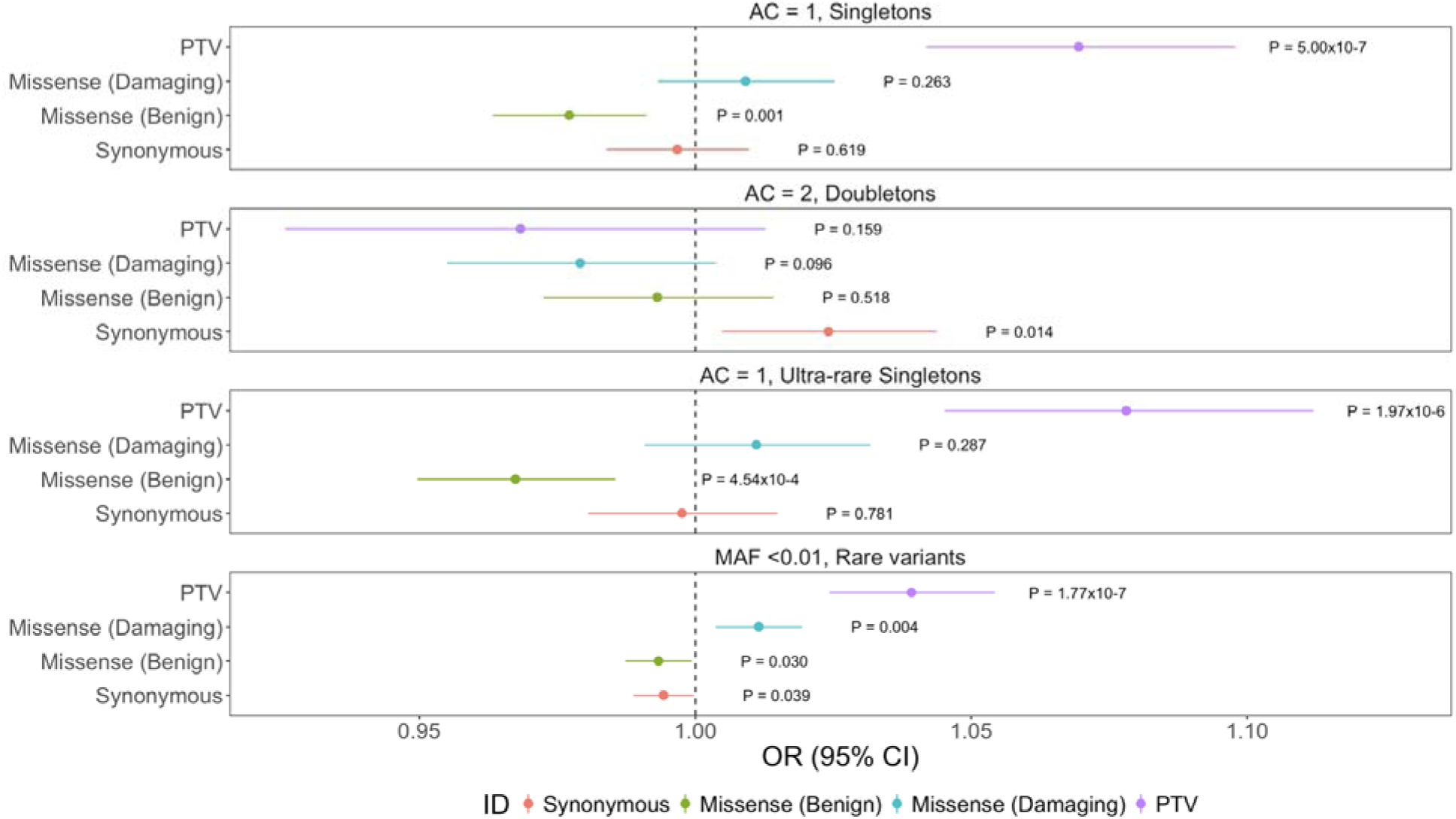
Exome wide enrichment of PTVs in ALS cases. Exome wide analysis of synonymous variants, benign missense variants, damaging missense variants, and PTVs within singletons (AC=1), doubletons (AC=2), ultra-rare singletons (AC=1, 0 in DiscovEHR), and rare variants (MAF<0.01 in our dataset, DiscovEHR and ExAC). Odds ratios and 95% confidence intervals for each class of variation are depicted by different colors.P-values are also displayed. Multiple test correction P-value: 0.0125.

When using model 4 where we restrict to ‘benign variation’ as the final covariate, the PTV signal is further enriched among singletons (OR: 1.12; 95% CI: 1.09-1.15; P-value: <2×10^-^ 16); ultra-rare singletons (OR: 1.10; 95% CI: 1.07-1.13; P-value: 1.53×10^-^10); and rare variants (OR: 1.04; 95% CI: 1.03-1.05; P-value: 1.47×10^-^7) (Figure S8). Interestingly, in this analysis, there is a consistent and a significant enrichment of damaging missense variants not observed in the previous analysis: singletons (OR: 1.06; 95% CI: 1.05-1.07; P-value: <2×10^-^16); ultra-rare singletons (OR: 1.03; 95% CI: 1.01-1.05; P-value: 6.33×10^-^5); and rare variants (OR: 1.01; 95% CI: 1.00-1.02; P-value: 3.24×10^-^3) (Figure S8).

In our analyses, we define PTVs as a frameshift variant, a splice acceptor variant, a splice donor variant, or a stop gained variant, which are due to insertions or deletions (indels), or single nucleotide variants (SNVs). We divided PTVs as either SNVs or indels given their differences in error rates (Lam et al., 2011; O’Rawe et al., 2013) and repeated the exome-wide analysis. The significant signal is present in both SNVs and indels: SNV singletons (OR: 1.05; 95% CI: 1.02-1.08; P-value: 2.99×10^-^3); indel singletons (OR: 1.10; 95% CI: 1.06-1.15; P-value: 5.75×10^-^6); SNV ultra-rare singletons (OR: 1.06; 95% CI: 1.02-1.10; P-value: 4.34×10^-^3); indel ultra-rare singletons (OR: 1.12; 95% CI: 1.06-1.17; P-value: 1.96×10^-^5); and SNV rare variants (OR: 1.03; 95% CI: 1.01-1.05; P-value: 6.48×10^-^4); indel rare variants (OR: 1.05; 95% CI: 1.03-1.07; P-value: 3.30×10^-^5) (Figure S10).

### Gene set testing: enrichment of rare variants in constrained genes

To determine whether we could concentrate the PTVs enrichment, we assessed multiple different gene sets. We evaluated (1) constrained genes under strong purifying selection (Lek et al., 2016; Samocha et al., 2014); (2) genes known to confer risk to ALS; (3) genes associated with clinically overlapping diseases; and (4) genes in which their expression is specific to the brain (“brain specific”).

Among constrained genes we observed a significant enrichment of singleton PTVs (OR: 1.23; 95% CI: 1.13-1.33; P-value: 7.74×10^-^7); ultra-rare singletons (OR: 1.27; 95% CI: 1.16-1.38; P-value: 5.76×10^-^8), and rare variants (OR: 1.33; 95% CI: 1.26-1.42; P-value: <2×10^-^16) (Figure 2A, Figure S11). We obtained similar results using model 4 (Figure S11). We did not observe a significant enrichment in damaging missense variants in constrained genes in model 3 except in rare variants: (OR: 1.08; 95% CI: 1.05-1.12; P-value: 4.15×10^-^6); however, there is enrichment using model 4: singletons (OR: 1.15; 95% CI: 1.09-1.20; P-value: 1.18×10^-^7); ultra-rare singletons (OR: 1.09; 95% CI: 1.03-1.16; P-value: 5.0×10^-^3), and rare variants (OR: 1.08; 95% CI: 1.04-1.11; P-value: 1.13×10^-^5) (Figure S11).

**Figure 2.**
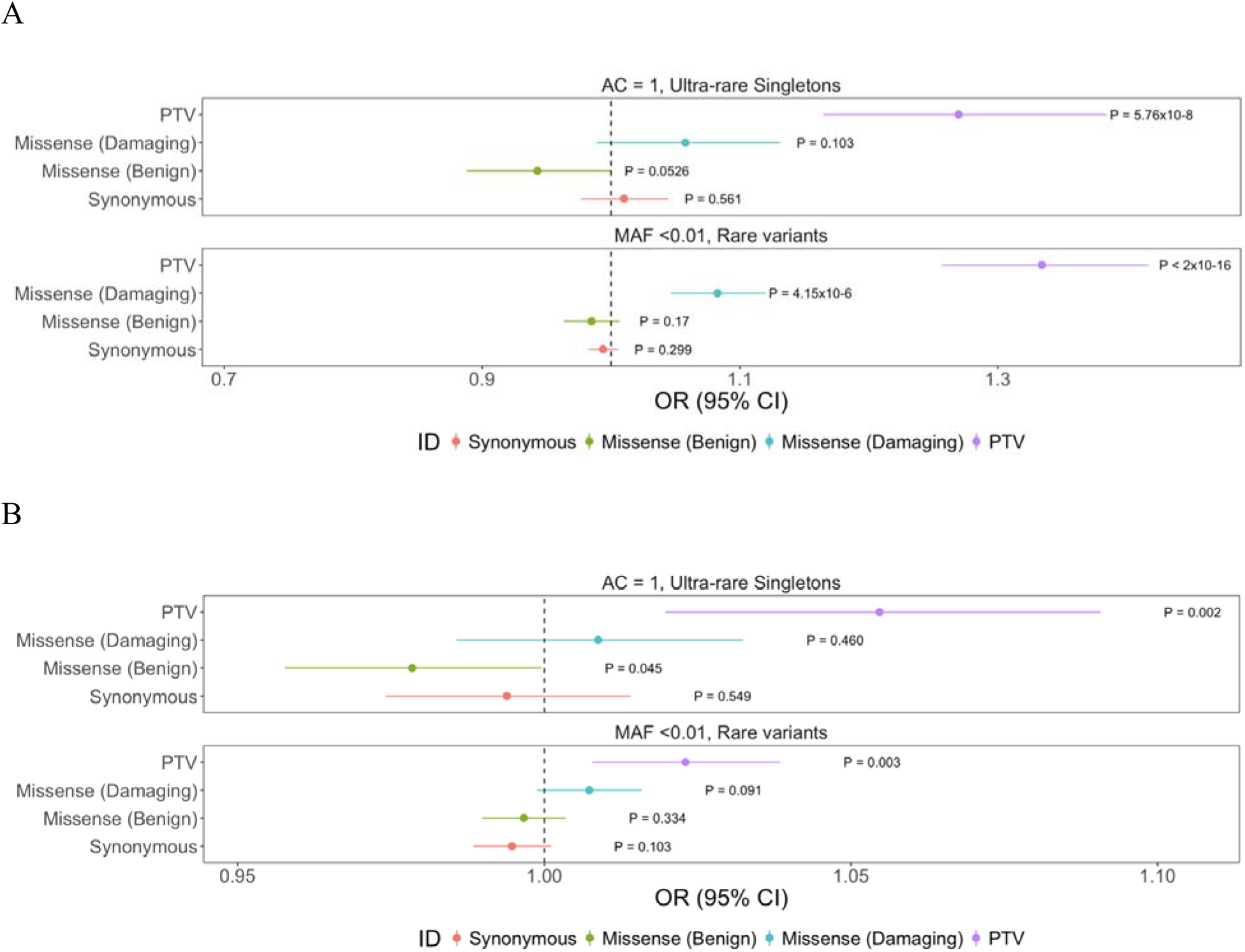
Enrichment of PTVs in constrained genes in ALS cases. (A) Analysis of constrained genes only in synonymous variants, benign missense variants, damaging missense variants, and PTVs within ultra-rare singletons (AC=1, 0 in DiscovEHR), and rare variants (MAF<0.01 in our dataset, DiscovEHR and ExAC). Odds ratios and 95% confidence intervals for each class of variation are depicted by different colors. P-values are also displayed. (B) Exome-wide analysis with constrained genes removed.

When we removed constrained genes, and repeated the exome wide analysis there is still a significant enrichment in singleton PTVs (OR: 1.05; 95% CI: 1.02-1.08; P-value: 3.30×10^-^4); ultra-rare singleton PTVs (OR: 1.05; 95% CI: 1.01-1.09; P-value: 1.96×10^-^3); and rare PTV variants (OR: 1.02; 95% CI: 1.00-1.04; P-value: 2.93×10^-^3) (Figure 2B, Figure S12). This was also observed in model 4 (Figure S12). In model 3, the significant enrichment in damaging missense variants in the rare variants category initially observed in constrained gene disappears: (OR: 1.00; 95% CI: 0.99-1.01; P-value: 0.091). In model 4, the signal remains with smaller effect sizes in singletons (OR: 1.07; 95% CI: 1.05-1.08; P-value: <2×10^-^16); in ultra-rare singletons (OR: 1.04; 95% CI: 1.02-1.06; P-value: 4.62×10^-^4), but not in rare variants (OR: 1.00; 95% CI: 0.99-1.01; P-value: 0.062) (Figure S12).

Next, we removed the potential effects of known ALS genes, and performed an exome-wide burden analysis. We did not remove the ALS genes *TBK1, NEK1, KIF5A, C21orf2, MOBP*, or *SCFD1* as these genes were discovered using datasets that contained a large subset of the same samples and can generate an amplified signal. The known ALS genes had negligible effects as the initial PTV signal remained without any changes in effect sizes: singletons (OR: 1.07; 95% CI: 1.04-1.10; P-value: 7.36×10^-^7); ultra-rare singletons (OR: 1.08; 95% CI: 1.04-1.11; P-value: 3.07×10^-^6); and rare variants (OR: 1.04; 95% CI: 1.02-1.06; P-value: 1.64×10^-^7) (Figure 3A, Figure S13). As expected, these observations were also supported by model 4 and the original signal in damaging missense variants persists (Figure S13). Importantly however, when including variants from *TBK1, NEK1, KIF5A, C21orf2, MOBP*, or *SCFD1*, the signals persist.

**Figure 3.**
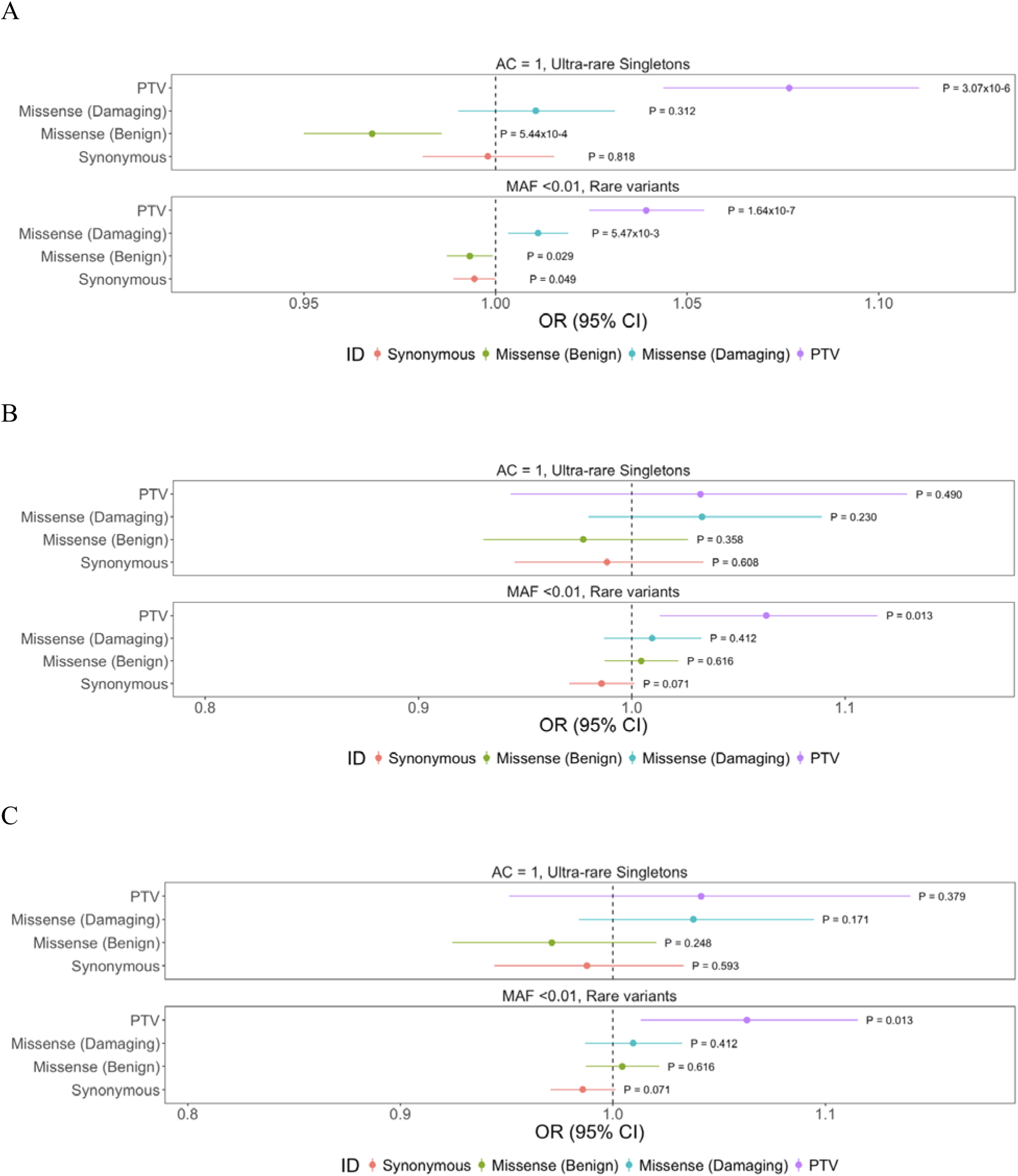
No enrichment of variants in known ALS genes, other related neurodegenerative disease genes, or brain specific genes. (A) Exome-wide analysis with known ALS genes removed in synonymous variants, benign missense variants, damaging missense variants, and PTVs within singletons (AC=1), doubletons (AC=2), ultra-rare singletons (AC=1, 0 in DiscovEHR), and rare variants (MAF<0.01 in our dataset, DiscovEHR and ExAC). Odds ratios and 95% confidence intervals for each class of variation are depicted by different colors. P-values are also displayed. (B) Analysis of other neurodegenerative disease genes (motor neuron diseases: primary lateral sclerosis, progressive muscular atrophy, progressive bulbar palsy, and spinal muscular atrophy; diseases with overlapping phenotypes: frontotemporal dementia, Parkinson’s disease, Pick’s disease, and Alzheimer’s disease). (C) Analysis of brain specific genes.

Although ALS is traditionally considered to be a disease of upper and lower motor neurons, more than 50% of ALS patients exhibit neuropsychological and cognitive deficits, with up to 30% of ALS patients meeting some diagnostic criteria for frontotemporal dementia, and some patients may also exhibit Parkinsonism or Parkinsonism-dementia (Aarsland et al., 2005; Farhan et al., 2017; Farhan et al., 2018; Hely et al., 2008; Strong et al., 2017; Swinnen and Robberecht, 2014). We surveyed the literature and tabulated a comprehensive list of genes associated with other motor neuron diseases such as primary lateral sclerosis, progressive muscular atrophy, progressive bulbar palsy, and spinal muscular atrophy. We also included genes associated with the following overlapping phenotypes: frontotemporal dementia, Parkinson’s disease, Pick’s disease, and Alzheimer’s disease. We did not observe a significant enrichment of variants in any class of variation, suggesting that the initial observation of PTV enrichment is likely not driven by genes associated with similar phenotypes (Figure 3B, Figure S15).

Finally, while the genes implicated in the development of ALS are not predominantly or preferentially expressed in motor neurons, nor are they all brain specific except for a select few, we tested whether there is a signal in brain specific genes as ALS is a neurodegenerative disease with the predominant symptoms affecting the central nervous system. We extracted a list of genes with specific brain expression generated using GTEx and performed the same burden analysis across classes of variation (Ganna et al., 2016). As expected, we did not observe any significant differences in PTVs or damaging missense variation in any allele frequency threshold (Figure 3C, Figure S16). These observations suggest the PTV signal initially observed in the exome-wide scan is also not driven by genes with specific brain expression, further corroborating the non-brain specific expression profiles of known ALS genes in the literature.

### Single gene burden analysis replicates previous ALS associations

To determine whether a single gene is enriched for variation in ALS cases (ALS-associated) or depleted in ALS cases (ALS-protective), we evaluated ultra-rare (AC=1, absent in DiscovEHR) and rare (MAF <0.001% in our dataset, DiscovEHR, and ExAC) PTVs and damaging missense variants. Within the ultra-rare variant category, no individual gene passed exome-wide significance. However, the top genes were known ALS genes: (1) *NEK1* (PTVs, OR: 12.21; 95% CI: 2.72-112.41; P-value: 7.32×10^-^5); (2) *OPTN* (PTVs, OR: 20.33; 95% CI: 2.89-878.93; P-value: 1.2×10^-^4); and (3) *SOD1* (dmis, OR: 46.91; 95% CI: 5.10-796.34; P-value: 5.03×10^-^6) (Figure S16). Within rare PTVs, only *NEK1* (OR: 12.8; 95% CI: 4.4-50.5; P-value: 4.59×10^-^9), passed exome-wide significance; the next top 9 most significant genes, which include *FUS*, a known ALS gene (OR: 26.4; 95% CI: 2.4-469.0; P-value: 1.29×10^-^3), are displayed in Table 1, Figure 4A; Figure S17A. Similarly, within damaging missense variants, *SOD1* (OR: 87.7; 95% CI: 10.6-1448.3; P-value: 7.5×10^-^11) was the only gene to pass exome-wide significance; the top 9 most significant genes are displayed in Table 2, Figure 4B; Figure S17B.

**Table 1.** Top hits in PTV model in initial (3,864 cases and 7,839 controls), and secondary datasets (3,864 cases and 28,910 controls)

| Gene | Initial OR | Initial P-value | Secondary OR | Secondary P-value |
| --- | --- | --- | --- | --- |
| *NEK1* | **12.8 (25, 4)** | **4.6x10<sup>-9</sup>\*** | **6.5 (25, 29)** | **3.0x10<sup>-10</sup><sup>#</sup>** |
| *DAK* | 42.7 (10, 0) | 1.5x10<sup>-5</sup> | 5.8 (10, 13) | 1.4x10<sup>-4</sup> |
| *IRAK3* | 11.2 (11, 0) | 2.0x10<sup>-4</sup> | 2.1 (11, 40) | 0.05 |
| *OPTN* | **6.6 (13, 4)** | **3.0x10<sup>-4</sup>** | **2.6 (13, 38)** | **6.9x10<sup>-3</sup>** |
| *SEC14L3* | 18.3 (9, 1) | 3.0x10<sup>-4</sup> | 1.4 (9, 48) | 0.31 |
| *VWA3B* | 5.7 (14, 5) | 3.0x10<sup>-4</sup> | 3.0 (14, 50) | 0.02 |
| *CDHR3* | 9.1 (9, 2) | 1.3x10<sup>-3</sup> | 1.4 (9, 70) | 0.9 |
| *ABCC2* | 3.1 (20, 13) | 1.3x10<sup>-3</sup> | 1.8 (20, 84) | 0.03 |
| *LRRC6* | 26.4 (6, 0) | 1.3x10<sup>-3</sup> | 1.2 (6, 38) | 0.64 |
| *FUS* | **26.4 (6, 0)** | **1.3x10<sup>-3</sup>** | **97.4 (6, 0)** | **2.7x10<sup>-6</sup><sup>#</sup>** |
| *GRIN3B* | 0.4 (0, 10) | 0.04 | 0.05 (0, 80) | 7.7x10<sup>-5</sup><sup>#</sup> |
| *HRCT1* | 0.2 (0, 15) | 4.1x10<sup>-3</sup> | 0.05 (0, 78) | 1.2x10<sup>-4</sup><sup>#</sup> |
| *IL3* | 14.2 (7, 1) | 2.4x10<sup>-3</sup> | 10.5 (7, 5) | 1.8x10<sup>-4</sup><sup>#</sup> |
| *DNAJC7* | 18.3 (4, 0) | 0.01 | 67.4 (4, 0) | 1.9x10<sup>-4</sup><sup>#</sup> |
| *KRT74* | 3.0 (9, 6) | 0.05 | 8.7 (9, 11) | 2.2x10<sup>-4</sup><sup>#</sup> |
| *SLC26A10* | 0.1 (0, 9) | 0.03 | 0.1 (0, 70) | 2.7x10<sup>-4</sup><sup>#</sup> |
| *FAM206A* | 4.1 (6, 3) | 0.07 | 11.2 (6, 4) | 3.7x10<sup>-4</sup><sup>#</sup> |
| ***TBK1*** | **22.3 (5, 0)** | **3.9x10<sup>-3</sup>** | **12.5 (5, 3)** | **9.3x10<sup>-4</sup><sup>#</sup>** |
| *KLHDC4* | 0.1 (0, 14) | 7.4 x10<sup>-3</sup> | 0.1 (0, 61) | 9.8x10<sup>-4</sup><sup>#</sup> |
| *DUOXA2* | 2.4 (7, 6) | 0.14 | 5.8 (7, 9) | 1.4x10<sup>-3</sup><sup>#</sup> |

\*Passed exome-wide significance (P-value <2.5x10<sup>-6</sup>) in first analysis (3,864 cases and 7,839 controls).
<sup>#</sup>OR direction is maintained in secondary analysis (3,864 cases and 28,910 controls) and P-value is lower.
Bolded genes have been previously reported in ALS.

**Table 2.**
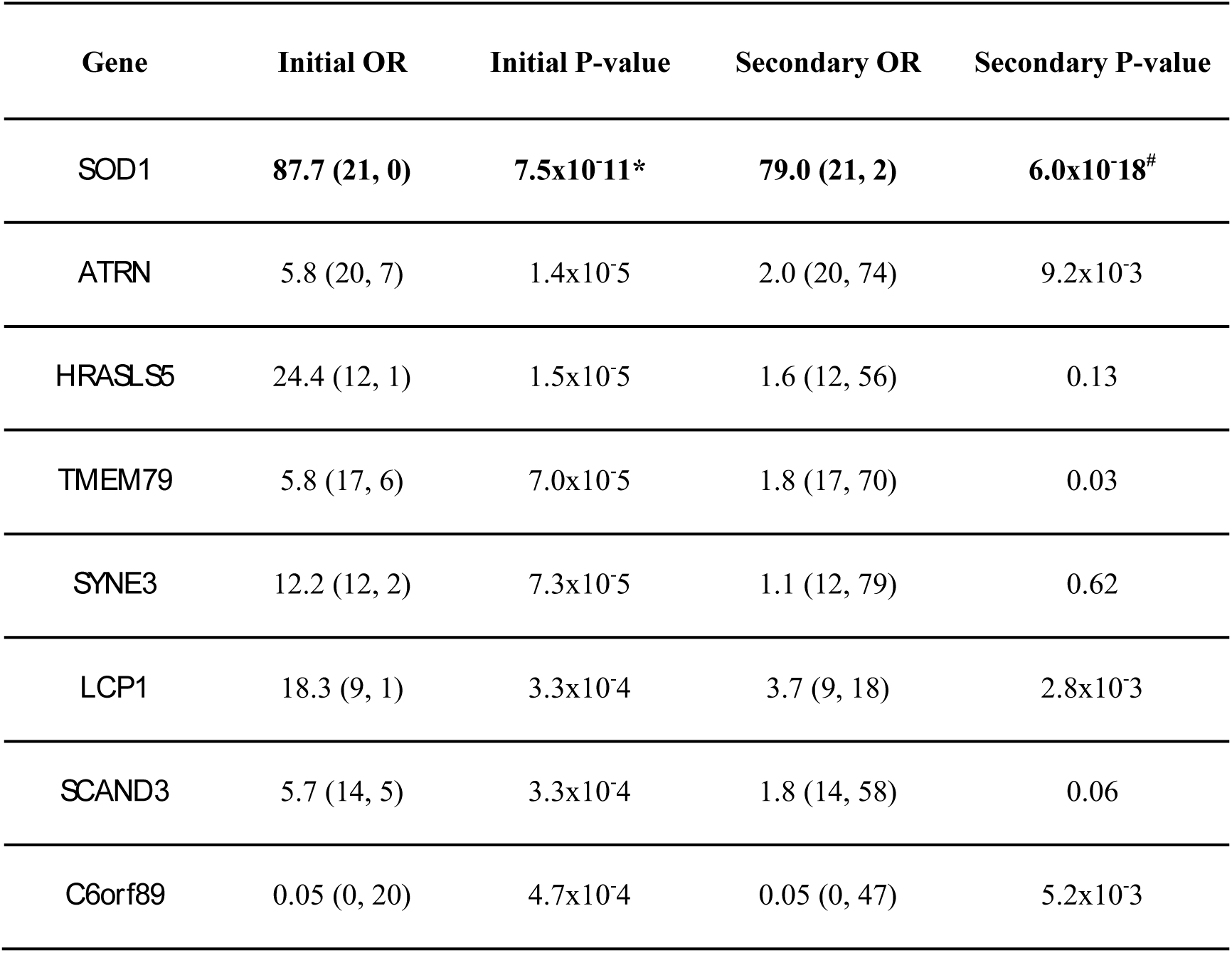
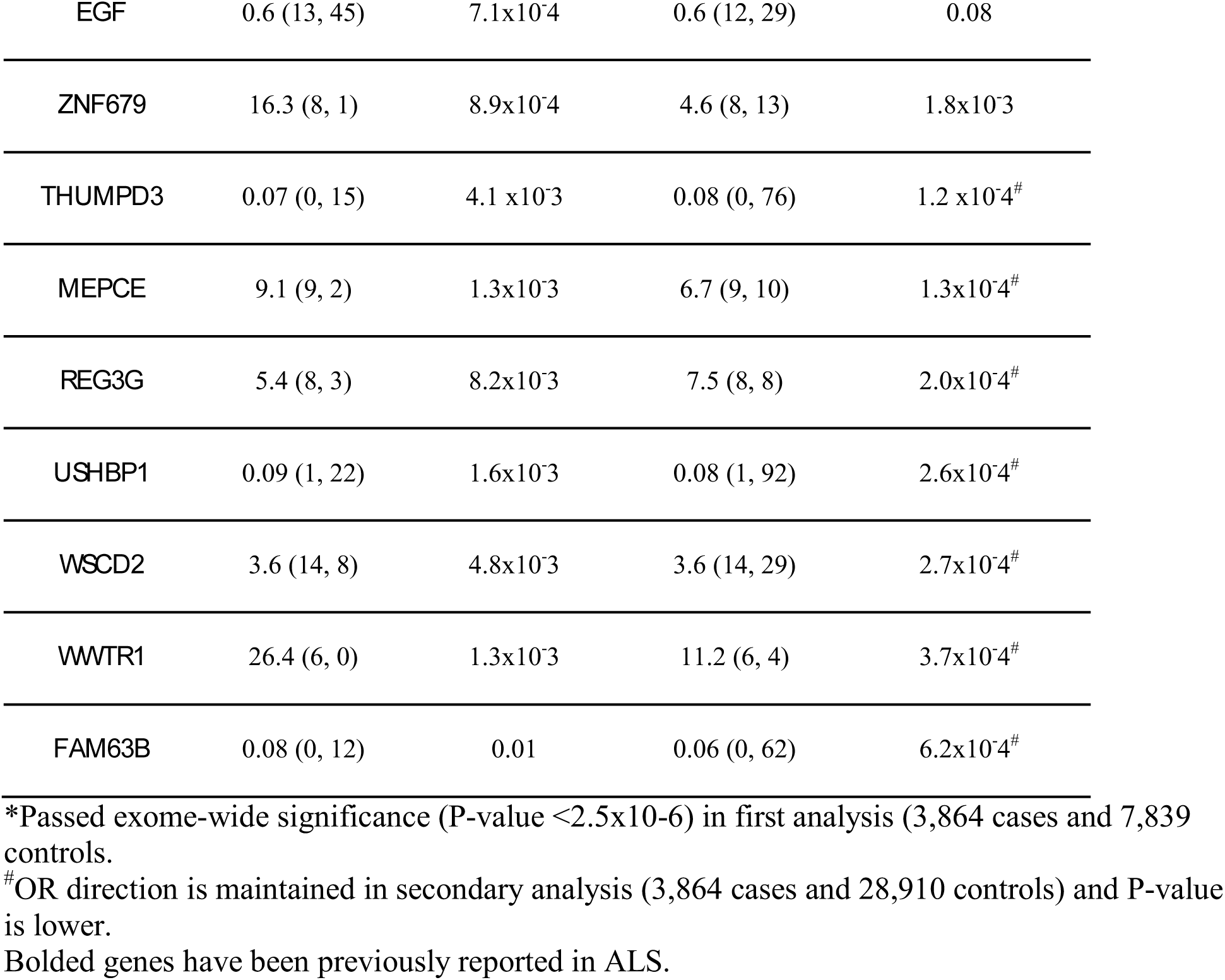
Top hits in damaging missense model in initial (3,864 cases and 7,839 controls), and secondary datasets (3,864 cases, 28,910 controls)

**Figure 4.**
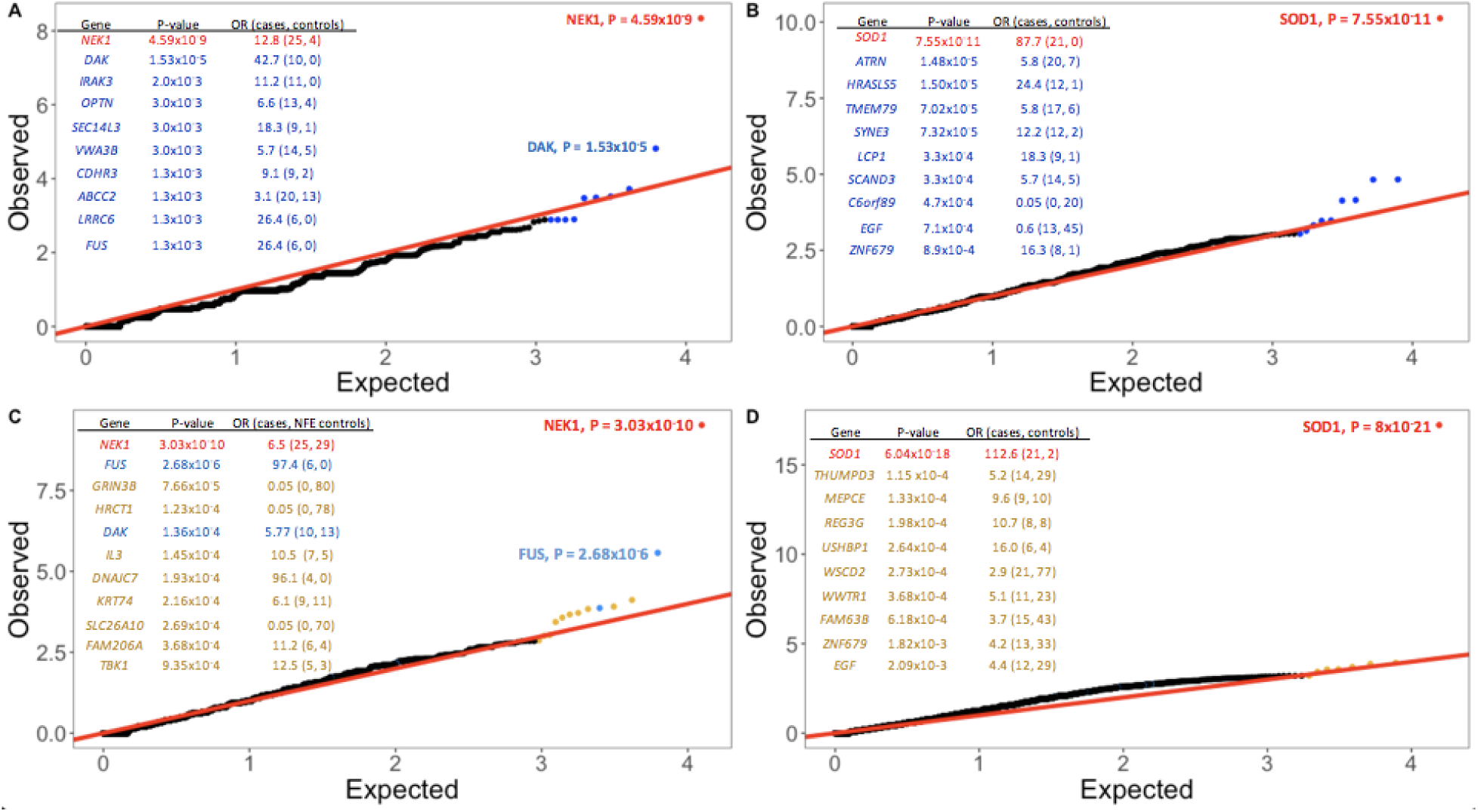
Quantile-quantile plot of discovery results for rare variants. (A) Rare PTVs (MAF<0.001% in dataset, DiscovEHR, and ExAC) in ALS dataset. PTVs in *NEK1* and *DAK* are enriched in ALS cases. The top 10 genes with their P-values are displayed. *NEK1* passes exome-wide significance. (B) Rare damaging missense variants (MAF<0.001% in dataset, DiscovEHR, and ExAC) in ALS dataset. Damaging missense variants in SOD1 are enriched in ALS cases. The top 10 genes with their P-values are displayed. *SOD1* passes exome-wide significance. (C) Rare PTVs (MAF<0.001% in dataset, DiscovEHR, and ExAC) in ALS cases with an additional 21,071 non-Finnish European controls from ExAC for a total of 28,910 controls. The top 10 genes with their P-values are displayed. *NEK1* is shown in red and passes exome-wide significance. Genes in blue were the most significant genes in the discovery analysis. FUS is shown in blue and approaches exome-wide significance. Genes in yellow were the most significant genes in the secondary analysis. (D) Rare damaging missense variants (MAF<0.001% in dataset, DiscovEHR, and ExAC) in ALS cases with an additional 21,071 non-Finnish European controls from ExAC for a total of 28,910 controls. The top 10 genes with their P-values are displayed. *SOD1* is shown in red and passes exome-wide significance. Genes in blue were the most significant genes in the discovery analysis. Genes in yellow were the most significant genes in the secondary analysis.

To replicate our findings, we included an additional 21,071 controls from ExAC that are of European descent (non-Finnish) and were not a part of any psychiatric or brain related studies, to eliminate any sample overlap. We performed the same burden analyses using 3,864 cases and 28,910 controls (7,839 controls within our dataset and 21,071 additional controls). In Tables 1 and 2, we display the most significant genes that were identified in the initial discovery and tabulate their OR and P-values for both the initial discovery cohort (3,864 cases and 7,839 controls) and the secondary analysis (3,864 cases and 28,910 controls). Within PTVs, *NEK1* is still the only gene that exceeds exome-wide significance (OR: 6.5; 95% CI: 3.6-11.5; P-value: 3.03×10^-^10) (Figure 4A, Figure S18). Of the next 9 most significant genes in the initial analysis, the only signal that was strengthened was in *FUS* (OR: 97.4; 95% CI: 8.8-1729.5; P-value: 2.68×10^-^6). Importantly, the signal in *OPTN*, a proposed ALS associated gene decreased (OR: 6.6; 95% CI: 2.0-27.8; P-value: 3.0×10^-^3 to OR: 2.6; 95% CI 1.25-4.93; P-value: 6.9×10^-^3). With the additional controls, multiple genes had similar ORs as the discovery analysis, with their respective P-values approaching significance (P-values ranging from 7.7×10^-^5-1.4×10^-^3). Most notably, the signal in *TBK1*, a proposed ALS gene based on Cirulli et al. strengthened: (initial analysis; OR: 22.3; 95% CI: 1.9-404.2; P-value: 3.9×10^-^3; secondary analysis: OR: 12.5; 95% CI: 2.4-80.3; P-value: 9.35×10^-^4). Furthermore, *DNAJC7*, which is a highly constrained gene (pLI = 1.0) had 4 PTVs in cases and 0 in controls in the discovery analysis (OR: 18.3; 95% CI:1.0 to 339.6; P-value: 0.01); and 0 PTVs in total controls, 28,910 (OR: 96.1; 95% CI: 4.9 to 1252.2; P-value: 1.9×10^-^4).

When considering the entire ExAC dataset (60,706 samples), there are 2 putative PTVs however, they are within multiallelic sites and are not sufficiently covered (17:40139856delG and 17:40142392insA). Similarly, within gnomAD (123,136 exomes and 15,496 genomes), there are 7 putative PTVs within the canonical transcript: 2 are within multiallelic sites (17:40139856delG and 17:40152552G>C); 1 is poorly called (17:40133978T>A); and there are 2 SNVs within the splice donor sites (17:40133871A>G and 17:40169357C>T), which may be rescued by neighboring sites with the same DNA sequence (17:40133867A and 17:40169349C). Nevertheless, there are still 2-4 putative PTVs in gnomAD, one of which is the same variant observed in 2 ALS cases within our dataset (17:40146902G>A, p.Arg156Ter), which may be pathological but incompletely penetrant. One important consideration is that gnomAD contains ALS exomes as well as exomes of other related neurodegenerative diseases or brain related phenotypes, and we are unable to detect whether *DNAJC7* PTVs are from neurodegenerative diseases exomes versus controls.

Within damaging missense variants, *SOD1* is still the only gene that exceeds exome-wide significance (OR: 79.0; 95% CI: 19.3-692.5; P-value: 6.0×10^-^18); however, the next 9 most significant genes no longer approach statistical significance. Similarly, when integrating additional controls, multiple genes approach significance (P-values ranging from 1.2×10^-^4-6.2×10^-^4) (Figure 4D, Figure S18).

## DISCUSSION

Herein, we have assembled the exomes of 3,864 ALS cases and 7,839 controls and observed an exome-wide enrichment of PTVs, which typically result in protein loss of function. The abundance of PTVs in ALS cases seems to be primarily driven by constrained genes, which are under strong purifying selection. When removing constrained genes, the initial exome-wide enrichment of PTVs remains; however, the effect sizes are much smaller, suggesting there are residual effects in other genes. Accordingly, we examined the effects of ALS associated genes by removing them from the exome-wide analysis and detected the same PTV enrichment initially observed. Importantly, a subset of cases was pre-screened for known pathogenic variants in a select number of known ALS genes and positive cases were eliminated prior to assembling the dataset, which will attenuate the estimates of effect size and significance for genes in this gene set.

Acknowledging the phenotypic variability of ALS, we also evaluated the effects of genes implicated in other motor neuron diseases such as primary lateral sclerosis, progressive muscular atrophy, progressive bulbar palsy, and spinal muscular atrophy; as well as genes associated with the following overlapping phenotypes: frontotemporal dementia, Parkinson’s disease, Pick’s disease, and Alzheimer’s disease. We did not observe a significant enrichment of variants in any class of variation, suggesting that the initial observation of excess PTVs do not reside in these genes. Lastly, the genes implicated in the development of ALS are not specifically expressed in motor neurons, nor are they brain specific, a perplexing observation given the specific degree of degeneration of upper and lower motor neurons. Nevertheless, we tested whether the signal in PTVs is concentrated in brain specific genes, a much larger gene set than ALS genes. As expected, we did not observe any significant enrichment in PTVs or damaging missense variation across brain specific genes. In summary, while much of the exome-wide signal in PTVs resides in constrained genes, it is likely there are other gene sets, not tested herein, that may partially explain the burden of PTVs in ALS cases.

The single gene burden analysis identified the most significant genes as *SOD1, NEK1*, and *FUS*, which are known ALS genes. No other individual gene passed exome-wide significance within our dataset (3,864 cases and 7,839 controls) and the additional controls in the secondary analysis (3,864 cases and 28,910 controls). Notably, in the secondary analysis, multiple genes with consistent OR and lower P-values than the initial analysis, surfaced. Within PTVs, these include: *GRIN3B, HRCT1, IL3*, and *DNAJC7*. Interestingly, both *GRIN3B* and *HRCT1* may offer protection against ALS: OR: 0.05, P-value: 7.7×10^-^5; OR: 0.05, P-value: 1.2×10^-^4, respectively; while *IL3* and *DNAJC7* may confer risk: OR: 10.5, P-value: 1.8×10^-^4; OR: 67.4, P-value: 1.9×10^-^4).

*DNAJC7* had 4 PTVs in 3,864 cases and 0 in 7,839 and 28,910 controls. When integrating the full ExAC (60,706 exomes) and gnomAD (123,136 exomes and 15,496 genomes) datasets, we observed 2 and 7 putative PTVs, respectively; however, the majority of the PTVs resided within poor quality or multiallelic sites. One caveat of using the full gnomAD dataset is that a subset of ALS exomes and other related neurodegenerative diseases have been deposited into this resource, which prevents us from delineating PTVs from ALS versus non-ALS exomes. When considering gnomAD genomes only (15,496 samples), there is only 1 putative PTV in *DNAJC7* due to a splice variant, which may be rescued by a neighboring nucleotide. According to the HPA RNA-seq normal tissues project (Fagerberg et al., 2014) and the Genotype-Tissue Expression (GTEx) project (Consortium, 2013), *DNAJC7* is ubiquitously expressed with elevated expression in the brain. *DNAJC7* encodes a molecular chaperone, DnaJ heat shock protein family (HSP40) member C7, and like all DNAJ proteins, contains an approximately 70 amino acid J-domain, which is critical for binding to HSP70 proteins (Jiang et al., 2007). There are approximately 50 DNAJ proteins, which are also classified as HSP40 proteins, that facilitate protein maintenance and quality control, such as folding of newly synthesized polypeptides, and clearance of degraded proteins (Kampinga and Craig, 2010; Lackie et al., 2017; Mayer and Bukau, 2005). Specifically, DNAJs act as co-chaperones for HSP70 proteins by regulating ATPase activity, aid in polypeptide binding, and prevention of premature polypeptide folding (Clerico et al., 2015; Mayer and Bukau, 2005).

Aberrant protein aggregation due to misfolding or accumulation of degraded proteins, is one of the pathological hallmarks of neurodegenerative diseases like Alzheimer’s disease, Parkinson’s disease, Huntington’s disease, prion disease, and ALS (Brundin et al., 2010; Imarisio et al., 2008; Irwin et al., 2013; Ross and Poirier, 2004; Uddin et al., 2018; Winklhofer et al., 2008). HSP proteins have a conserved and central role in protein function by aiding in their folding and stabilization, and the clearance of misfolded proteins, ultimately diminishing protein aggregates and the associated pathologies. However, cellular stress such as exposure to environmental toxins, fluctuations in temperature, chemical stress, cell injury, or aging, can influence the dynamics of the protein quality control network allowing misfolded proteins to go undetected thereby triggering neurotoxicity (Gidalevitz et al., 2006; Voisine et al., 2010). Furthermore, abnormal expression of HSP70 and DNAJ genes leads to the formation of protein aggregates in models of Alzheimer’s disease (Brehme et al., 2014), Parkinson’s disease (Auluck et al., 2002; Roodveldt et al., 2009), Huntington’s disease (Brehme et al., 2014; Wacker et al., 2009), prion disease (Jones et al., 2004; Kovacs et al., 2001), and ALS (Chen et al., 2016; Udan-Johns et al., 2014; Zhang et al., 2010). In light of these studies, elevated HSP expression is thought to be beneficial in preventing or in halting neurodegenerative disease progression (Benatar et al., 2018). For example, overexpression of *DNAJB6b* and *DNAJB8* suppressed toxic protein aggregation (Hageman et al., 2010); while overexpression of HSP70 in neuroglioma cells decreased the formation of alpha-synuclein fibrils (Outeiro et al., 2008). Within ALS models, overexpression of HSPB8 promoted clearance of mutant SOD1 (Crippa et al., 2010); double transgenic mice overexpressing HSP27 and mutated SOD1 exhibited increased survival of spinal motor neurons than mice overexpressing a *SOD1* mutation only, however, the neuroprotective effects were not sustained in later stages of the disease (Sharp et al., 2008). Finally, *DNAJB2*, which when mutated can cause autosomal recessive spinal muscular atrophy, was overexpressed in mice motor neurons also expressing a *SOD1* mutation (p.Gly93Ala), and led to reduced mutant SOD1 aggregation and improved motor neuron survival (Novoselov et al., 2013).

While no novel genes that pass exome-wide significance have been unveiled herein, the significant exome-wide enrichment of PTVs, which seem to primarily reside in constrained genes, substantiates the polygenicicity of ALS (McLaughlin et al., 2017; van Rheenen et al., 2016). Therefore, the integration of a larger number of ALS samples can potentially identify novel genes that are highly constrained, like *DNAJC7*, that may confer risk to ALS.

## EXPERIMENTAL PROCEDURES

### Study overview

The familial ALS (FALS) and the ALS Genetics (ALSGENS) consortia were assembled to aggregate the existing ALS sequencing data in the community to improve the power to discover novel genetic risk factors for ALS. Herein, we describe our approach of assembling the largest ALS exome case-control study to date.

### Sample acquisition

Blood samples were collected from subjects following appropriate and informed consent in accordance with the Research Ethics Board at each respective recruiting site within the CReATe, FALS, and ALSGENS consortia. All samples known to be carriers of the *C9orf72* hexanucleotide expansion (G4C2) were excluded from the study. Additionally, prior to exome sequencing, a subset of the samples (approximately 2,000) were genotyped and screened for known variants in known ALS genes, *SOD1, FUS*, and *TARDBP*; and were only included in our study if they were found to be negative for the variants tested.

Exome sequencing data for control samples were downloaded from dbGAP and were not enriched for (but not specifically screened for) ALS or other neurodegenerative disorders. Control samples were matched to case samples with respect to similar capture kits and coverage levels. The age of control samples was not provided for all samples but in general, controls were older than typical age of onset of ALS.

### Whole exome sequencing

15,722 DNA samples were sequenced at the Broad Institute, Guy’s Hospital, McGill University, Stanford University, HudsonAlpha, and University of Massachusetts, Worcester. Samples were sequenced using the exome Agilent All Exon (37MB, 50MB, or 65MB), Nimblegen SeqCap EZ V2.0 or 3.0 Exome Enrichment kit, Illumina GAIIx, HiSeq 2000, or HiSeq 2500 sequencers according to standard protocols.

All samples were joint called together and were aligned to the consensus human genome sequence build GRCh37/hg19; and BAM files were processed using BWA Picard. Genotype calling was performed using the Genome Analysis Toolkit’s (GATK) HaplotypeCaller and was performed at the Broad Institute as previously described (Ganna et al., 2017, BioRxiv, doi: https://doi.org/10.1101/148247).

### Hail software and quality control

We used Hail, an open-source, scalable framework for exploring and analyzing genomic data https://hail.is/ to process the data. All quality control steps were performed using Hail 0.1 (Table S1).

#### (1) Sample QC and Variant QC

Samples with high proportion of chimeric reads (>5%) and high contamination (>5%) were excluded. Samples with poor call rates (<90%), mean depth <10x, or mean genotype-quality <65 were also eliminated from further analysis.

For variant QC, we restricted to GENCODE coding regions, independent of capture interval, where both Agilent and Illumina exomes surpass 10x mean coverage. We restricted to ‘PASS’ variants in GATK’s Variant Quality Score Recalibration (VQSR) filter. Individual genotypes were filtered (set to missing) if they did not meet the following criteria: 1) genotype depth (g.DP) 10 or greater 2) Allele balance >=0.2 in heterozygous sites or <= 0.8 for homozygous reference and homozygous alternate variants 3) Genotype quality (GQ)> 20. Finally, we selected variants with call rate >90% and Hardy-Weinberg equilibrium test P-value >1×10^-^6. For quality control analysis, see Table S1 and Figures S1.

#### (2) Sex imputation

We used the X chromosome inbreeding coefficient to impute sample sex. Samples with an X chromosome inbreeding coefficient >0.8 were classified as males and samples with an X chromosome inbreeding coefficient <0.4 were classified as females. Samples within <0.8 and >0.4 were classified as having ambiguous sex status, and therefore were excluded from the dataset (Figure S2, Table S1).

#### (3) Principal component analysis

Principal component analysis (PCA) was performed using Hail. We used a subset of high confidence SNPs in the exome capture region to calculate the principal components (Figure S3A). We used only ancestry-matched cases and controls as indicated by overlapping population structure (Figure S3B-3D). Furthermore, we used 1000 Genomes samples to determine the general ethnicity of the ALS dataset. The majority of the samples in the ALS dataset were reported to be of European descent and this was confirmed by PCA with 1000 Genomes samples (Figure S4, Table S1).

#### (4) Relatedness check

We included only unrelated individuals (IBD proportion < 0.2) (Figure S5, Table S1).

#### (5) Variant annotation

We annotated protein-coding variants into four classes: (1) synonymous; (2) benign missense;(3) damaging missense; and (4) protein-truncating variants (PTV). Using VEP annotations (Version 85) (McLaren et al., 2016), we classified synonymous variants as: “synonymous_variant”, “stop_retained_variant”, and “incomplete_terminal_codon_variant”. Missense variants were classified as: “inframe_deletion”, “inframe_insertion”, “missense_variant”, “stop_lost”, “start_lost”, and “protein_altering_variant”. Furthermore, benign missense variants were predicted as “tolerated” and “benign” by PolyPhen-2 and SIFT, respectively; whereas damaging missense variants were predicted as “probably damaging” and “deleterious”. Finally, PTVs were classified as: “frameshift_variant”, “splice_acceptor_variant”, “splice_donor_variant”, and “stop_gained”.

#### (6) Allele frequency categorization

Allele frequencies were estimated within our case-control sample, and from two external exome sequence databases, DiscovEHR and ExAC (Lek et al., 2016). DiscovEHR is a publicly available database with >50,000 exomes of ostensibly healthy participants. ExAC is a mixture of healthy controls and complex disease patients, and we restricted to the non-psychiatric subset of ExAC for allele frequency estimation. Of note, many of our controls are present in the ExAC database, so we restricted to the DiscovEHR cohort to determine ultra-rare singletons. We did not use gnomAD for this analysis as our cases and our controls have been deposited into this resource.

We classified variant allele frequency using the following criteria: (1) singletons, which are variants present in a single individual in our dataset (allele count, AC =1); (2) doubletons, which are present in two individuals in our dataset (AC = 2); (3) ultra-rare singletons, which are singletons in our dataset and are absent in DiscovEHR (AC = 1, 0 in DiscovEHR); and finally, (4) rare variants, which have a MAF of <0.01% in our dataset (11,703 samples), in ExAC (non-psychiatric studies, >45,000 samples) and in DiscovEHR (>50,000 samples).

### Multivariate models used for analysis

To determine whether an enrichment of a specific class of variation was present in ALS cases versus controls, we ran multiple Firth logistic regression models. The Firth penalization is used in the likelihood model due to the low counts in many tests, and helps to minimize the type I error rate when multiple covariates are included in the model (Wang, 2014). Model 1 predicted ALS case-control status solely from variant count (Figure S6); Model 2 incorporated multiple covariates: (1) sample sex, (2) sample population structure from the first 10 principal components (Figure S7); Model 3 incorporated all covariates used the second model along with (3) sample total exome count, which is the exome-wide count of variants in the specific frequency class tested. Finally, Model 4 is similar to Model 3, but instead uses the “benign variant” count as a covariate, which is the exome-wide count of synonymous variants and benign missense variants only, rather than total exome count (Figure S8). Model 3, which we considered to be the most conservative model to represent the dataset, was used as the preferred model for our analysis (Figure S9).

### Exome-wide burden

The four Firth logistic regression models above were used to predict case-control status from exome-wide counts of synonymous, missense, and PTV variants. Given that sequencing errors are more prevalent when calling insertions or deletions (indels), we divided variants within the PTV category as either 1) SNV-based PTVs or 2) indel-based PTVs, due to single nucleotide variants (SNVs) or indels, respectively. This ensures that any enrichment observed in PTVs is not solely from indel-based PTVs.

### Gene sets

#### (1) Constrained genes (pLI genes: 3,488, constrained missense genes: 1,730)

We evaluated whether variation in loss of function intolerant (pLI) genes are associated with ALS using the same approach as described in the exome-wide approach however, we extracted only high pLI genes from the exome. We obtained the genic pLI intolerance metrics from Lek et al., 2016 available online:

(ftp://ftp.broadinstitute.org/pub/ExAC_release/release0.3/functional_gene_constraint/). For PTVs, we used genes with a probability of loss-of-function intolerant (pLI) >0.9. We also evaluated missense constrained genes generated by Samocha et al., 2014. For missense variants, we used genes with a z-score of >3.09.

#### (2) ALS associated genes (38 genes)

We also examined exome-wide burden with known ALS genes removed. The list of ALS genes are as follows: *TARDBP, DCTN1, ALS2, CHMP2B, ARHGEF28, MATR3, SQSTM1, FIG4, HNRNPA2B1, C9orf72, SIGMAR1, VCP, SETX, OPTN, PRPH, HNRNPA1, DAO, ATXN2, ANG, FUS, PFN1, CENPV, TAF15, GRN, MAPT, PNPLA6, UNC13A, VAPB, SOD1, NEFH, ARPP21*, and *UBQLN2*. We did not remove *TBK1, NEK1, KIF5A, C21orf2, MOBP*, or *SCFD1* as these genes were discovered using datasets that contained a large subset of the same samples. We also performed an exome-wide burden analysis with all proposed ALS genes removed.

#### (3) Neurodegenerative disease genes (120 genes)

We investigated whether genes associated with other neurodegenerative phenotypes showed enrichment in ALS cases. We included the following motor neuron diseases: primary lateral sclerosis, progressive muscular atrophy, progressive bulbar palsy, and spinal muscular atrophy. We also used genes associated with Parkinson’s disease, frontotemporal dementia, Pick’s disease, and Alzheimer’s disease as patients with ALS can also present with frontotemporal dementia, cognitive impairment, or Parkinsonism (Table S2).

#### (4) Brain expressed genes (2,650 genes)

We evaluated whether genes expressed specifically in the brain were enriched for variation in our dataset. For this analysis, we used brain specific genes generated by Ganna et al., 2016.

### Single gene burden analysis

#### (1) ALS dataset (3,864 cases and 7,839 controls)

To determine whether a single gene is enriched or depleted for rare protein-coding variation in ALS cases, we performed a burden analysis using Fisher’s exact test as well as SKAT, with previously defined covariates (sample sex, PC1-PC10, and total exome count). Exome-wide correction for multiple testing was set at (P<2.5×10^-^6), which was the 5% type-I error rate multiplied by the number of genes tested. We performed four different tests in ALS cases and controls: (1) ultra-rare PTVs (AC=1 and absent in DiscovEHR); (2) ultra-rare damaging missense variants (AC=1 and absent in DiscovEHR); (3) rare PTVs (MAF <0.001% in the dataset, DiscovEHR, and ExAC); and (4) rare damaging missense variants (MAF <0.001% in the dataset, DiscovEHR, and ExAC).

#### (2) ALS dataset and additional controls (3,864 cases and 28,910 controls)

We also included an additional 21,071 samples from ExAC that are of European descent (non-Finnish) and were not a part of any psychiatric or brain related studies, to eliminate any sample overlap. Furthermore, to mitigate against false discoveries, in addition to passing our QC filters, we ensured each variant also passed gnomAD (123,136 exomes and 15,496 genomes) QC filters. We included variants that were either a singleton (AC=1) in gnomAD or completely absent to ensure we minimize the inclusion of an excess of variants that passed gnomAD QC, that were rare (MAF <0.001%), yet were still observed in a very high number of individuals and were likely, false positive variants. The additional 21,071 samples allowed us to perform in part, a replication (secondary) analysis of the genes that approached statistical significance (P<2.5×10^-^6) and determine whether their OR and P-values are maintained and exceed statistical significance, respectively. Additionally, we also used the 21,071 controls to increase statistical power to detect any gene discoveries not detected in the original dataset. Importantly, we did not perform a joint PCA on the 21,071 non-Finnish European controls and our dataset, therefore, we are unable to completely match the ancestry of our dataset.

## SUPPLEMENTAL INFORMATION

Supplemental Information includes five tables and eighteen figures.

## CONSORTIA

The members of the Clinical Research in ALS and Related Disorders for Therapeutic Development (CReATe) Consortium are Michael Benatar, J. Paul Taylor, Gang Wu, Evadnie Rampersaud, Joanne Wuu, Rosa Rademakers, Stephan Zuchner, Rebecca Schule, Jacob McCauley, Sumaira Hussain, Anne Cooley, Marielle Wallace, Christine Clayman, Richard Barohn, Jeffrey Statland, John Ravits, Andrea Swenson, Carlayne Jackson, Jaya Trivedi, Shaida Khan, Jonathan Katz, Liberty Jenkins, Ted Burns, Kelly Gwathmey, James Caress, Corey McMillan, Lauren Elman, Erik Pioro, Jeannine Heckmann, Yuen So, David Walk, Samuel Maiser, and Jinghui Zhang.

The members of the FALS Consortium are Giuseppe Lauria, Orla Hardiman, Russell L McLaughlin, Letizia Mazzini, Stefano Duga, Anneloor L M A ten Asbroek, Frank Baas, Lucia Corrado, Sandra D’Alfonso, Jonathan D, Glass, Meraida Polak, Seneshaw Asress, Antonia Ratti, Cinzia Tiloca, Claudia Colombrita, Daniela Calini, Federico Verde, Nicola Ticozzi, Vincenzo Silani, Claudia Fallini, Diane McKenna-Yasek, John E. Landers, Kevin P. Kenna, Pamela Keagle, Peter C. Sapp, Robert H. Brown Jr, Cinzia Bertolin, Gianni Sorarù, Giorgia Querin, Giacomo P. Comi, Roberto Del Bo, Stefania Corti, Cristina Cereda, Mauro Ceroni, Stella Gagliardi, Garth A. Nicholson, Ian P. Blair, Kelly L. Williams, Karen E. Morrison, Hardev Pall, Ammar Al-Chalabi, Andrew King, Athina Soragia Gkazi, Bradley N. Smith, Caroline Vance, Christopher E. Shaw, Claire Troakes, Jack W. Miller, Safa Al-Sarraj, Simon D. Topp, Michael Simpson, Nick W. Parkin, Claire S. Leblond, Guy A. Rouleau, Patrick A. Dion, Jacqueline de Belleroche, Kevin Talbot, Martin R. Turner, Pamela J. Shaw, P. Nigel Leigh, Alberto García-Redondo, Jesús Esteban-Púrez, José Luis Muñoz-Blanco, Barbara Castellotti, Cinzia Gellera, Franco Taroni, Viviana Pensato.

The members of the ALS Sequencing Consortium are Andrew S. Allen, Stanley Appel, Robert H. Baloh, Richard S. Bedlack, Braden E. Boone, Robert Brown, John P. Carulli, Alessandra Chesi, Wendy K. Chung, Elizabeth T. Cirulli, Gregory M. Cooper, Julien Couthouis, Aaron G. Day-Williams, Patrick A. Dion, Summer Gibson, Aaron D. Gitler, Jonathan D. Glass, David B. Goldstein, Yujun Han, Matthew B. Harms, Tim Harris, Sebastian D. Hayes, Angela L. Jones, Jonathan Keebler, Brian J. Krueger, Brittany N. Lasseigne, Shawn E. Levy, Yi-Fan Lu, Tom Maniatis, Diane McKenna-Yasek, Timothy M. Miller, Richard M. Myers, Slave Petrovski, Stefan M. Pulst, Alya R. Raphael, John M. Ravits, Zhong Ren, Guy A. Rouleau, Peter C. Sapp, Neil A. Shneider, Ericka Simpson, Katherine B. Sims, John F. Staropoli, Lindsay L. Waite, Quanli Wang, Jack R. Wimbish, and Winnie W. Xin.

## ACKNOWLEDGEMENTS

We thank and acknowledge the consent and cooperation of all study participants. Many thanks to Felecia Cerrato for helping us assemble the dataset and providing general project management; and to Tim Poterba, Dr. Jon Bloom, Daniel King, and Dr. Cotton Seed for their assistance in Hail. We also thank Katariina Mamia for her help in sample preparation. The CReATe consortium (U54NS092091) is part of Rare Diseases Clinical Research Network (RDCRN), an initiative of the Office of Rare Diseases Research (ORDR), NCATS. This consortium is funded through collaboration between NCATS, and the NINDS. Additional support is provided by the ALS Association (17-LGCA-331). SMKF is supported by the ALS Canada Tim E. Noël Postdoctoral Fellowship.

## AUTHOR CONTRIBUTIONS

SMKF, MJD, and BMN conceived and designed the experiments. SMKF, HP, BNS, ST, ER, GW, JW, AG, CReATe Consortium, FALS Consortium, ALSGENS Consortium, KE, RR, JLM, RS, SZ, MB, JPT, MN, BT, CES, DBG, MBH, and BMN collected samples, prepared samples for analysis, or were involved in clinical evaluation. MB and JPT were the lead contacts for the CReATe Consortium. SDT and CES were the lead contacts for the FALS Consortium. DBG and MBH were the lead contacts for the ALSGENS Consortium. SMKF performed all experiments and executed data analyses. DPH, LEA, AEB, and SDT provided analysis suggestions. SMKF performed the primary writing of the manuscript with guidance from DPH, CC, MJD, and BMN. All authors approved the final manuscript. MJD and BMN supervised the research.

## DECLARATION OF INTERESTS

MAN participation is supported by a consulting contract between Data Tecnica International and the National Institute on Aging, NIH, Bethesda, MD, USA, as a possible conflict of interest Dr. Nalls also consults for Lysosomal Therapeutics Inc, the Michael J. Fox Foundation and Vivid Genomics among others.

